# *N*-linked glycosylation of the M-protein variable region: Glycoproteogenomics reveals a new layer of personalized complexity in multiple myeloma

**DOI:** 10.1101/2023.04.05.535540

**Authors:** Pieter Langerhorst, Melissa Baerenfaenger, Purva Kulkarni, Simon Nadal, Charissa Wijnands, Merel A. Post, Somayya Noori, Martijn M. vanDuijn, Irma Joosten, Thomas Dejoie, Alain J. van Gool, Jolein Gloerich, Dirk J. Lefeber, Hans J.C.T. Wessels, Joannes F.M. Jacobs

**Affiliations:** Department of laboratory Medicine, Radboud University Medical Center, Nijmegen, The Netherlands; Department of Neurology, Donders Institute for Brain, Cognition, and Behavior, Radboud University Medical Center, Nijmegen, Netherlands; Division of BioAnalytical Chemistry, Vrije Universiteit Amsterdam, Amsterdam, The Netherlands; Medical BioSciences Department, Radboud University Medical Center, Nijmegen, The Netherlands; Department of Human Genetics, Radboud University Medical Center, Nijmegen, The Netherlands; CY Cergy Paris Université, CNRS, BioCIS, Cergy-Pontoise, France; Department of Neurology, Erasmus University Medical Center, Rotterdam, the Netherlands; Biochemistry Laboratory, Centre Hospitalier Universitaire (CHU), Nantes, France

**Author notes:** these authors contributed equally (Hans J.C.T. Wessels, Joannes F.M. Jacobs). Corresponding author; Department of Laboratory Medicine, Radboud University Medical Center, Laboratory Medical Immunology (Route 469), Geert Grooteplein 10, 6525 GA Nijmegen, the Netherlands.

## Abstract

Multiple Myeloma (MM) is a plasma cell malignancy characterized by a monoclonal expansion of plasma cells that secrete a characteristic M-protein. This M-protein is crucial for diagnosis and monitoring of MM in the blood of patients. Recent evidence has emerged suggesting that N-glycosylation of the M-protein variable (Fab) region contributes to M-protein pathogenicity, and that it is a risk factor for disease progression of plasma cell disorders. Current methodologies lack the specificity to provide a site-specific glycoprofile of the Fab regions of M-proteins. Here, we introduce a novel glycoproteogenomics method that allows detailed M-protein glycoprofiling by integrating patient specific Fab region sequences (genomics) with glycoprofiling by glycoproteomics. Genomic analysis uncovered a more than two-fold increase in the Fab Light Chain N-glycosylation of M-proteins of patients with Multiple Myeloma compared to Fab Light Chain N-glycosylation of polyclonal antibodies from healthy individuals. Subsequent glycoproteogenomics analysis of 41 patients enrolled in the IFM 2009 clinical trial revealed that the majority of the Fab N-glycosylation sites were fully occupied with complex type glycans, distinguishable from Fc region glycans due to high levels of sialylation, fucosylation and bisecting structures. Together, glycoproteogenomics is a powerful tool to study *de novo* Fab N-glycosylation in plasma cell dyscrasias.

## Introduction

Multiple Myeloma (MM) is a plasma cell malignancy characterized by a monoclonal expansion of plasma cells in the bone marrow. These malignant plasma cells produce and secrete a characteristic monoclonal immunoglobulin (M-protein)^1^. M-protein diagnostics is imperative for diagnosis and monitoring of patients with MM^2^. Similar to other antibodies, the M-protein consists of a constant (Fc) and a variable (Fab) region. The Fc region is immunoglobulin (Ig) type specific and harbors the effector functions of the antibody, and the Fab region defines the binding specificity of the antibody and is both patient- and myeloma specific^3^. The variable Fab domain arises through unique rearrangements of germline V(D)J genes and subsequent somatic hypermutations^4^. These unique regions and their derived clonotypic peptides can be used for blood-based monitoring of MM minimal residual disease activity^5–8^.

Protein N-glycosylation is an important post-translational modification (PTM), where oligosaccharides are enzymatically attached to asparagine residues in proteins^9–11^. Immunoglobulins are the most abundant glycoproteins in the blood. All immunoglobulins (Ig) contain glycans that are linked to conserved motifs in the Fc-region. This Ig glycosylation is known to affect Ig function in Fc Receptor and complement component 1q binding^12,13^. In addition, the Fab region may also acquire N-linked glycans. This, however, requires the introduction of amino acid consensus motifs for N-glycosylation during somatic hypermutation^12^. The estimated incidence rate of Fab N-glycosylation is ~15%, yet the exact role of Fab N-glycans is not yet established^13^. Recent studies suggest that Fab glycans are involved in immune regulation, thereby playing a role in autoimmunity and also in the clinical effect of intravenous Ig products^12–15^.

Similar to polyclonal antibodies, an M-protein Fab region may also be glycosylated. Recently, evidence has emerged that the presence of M-protein light chain (LC) Fab N-glycosylation contributes to the M-protein pathogenicity in amyloid light chain(AL)-amyloidosis^16^. Additionally, M-protein Fab-glycosylation of the LC is reported as a risk factor for Monoclonal Gammopathy of Undetermined Significance (MGUS) disease progression to AL-amyloidosis, MM and other plasma cell dyscrasias^17^.

Methods to study Fab N-glycosylation have classically been based on glycan binding lectins or released glycan analysis by Mass Spectrometry (MS) of enriched Fab regions of IgG’s^12–14,18,19^. Recently, direct analysis of intact M-protein light chains by MALDI-TOF MS has been proposed to detect mass shifts corresponding with glycan mass distances^16^. However, all of the methods mentioned above cannot resolve the M-protein glycosylation status at the level of individual N-glycosylation sites which is required to gain novel insights into M-protein glycobiology. Moreover, since N-glycosylation is a highly heterogenous PTM with variable glycan structures and occupancy rates, we believe that site-specific glycosylation data is key to discern the biological or clinical consequences of Fab N-glycosylation^20^. We hypothesize that the unique M-protein Fab N-glycosylation status can be characterized by personalized genomic analysis of the myeloma plasma cells to derive reference M-protein sequences to enable subsequent glycoproteomics for detailed site-specific glycosylation analysis^21^. This may expand our knowledge on M-protein pathogenicity.

Here, we introduce a novel glycoproteogenomics method, through integration of genomics and glycoproteomics, to predict and confirm *de novo* Fab N-glycosylation profiles of M-proteins from patients with MM in a site-specific manner. Based on genomic data, we showed that M-proteins from patients with MM more often have a Fab N-glycosylation site compared to Ig from healthy individuals. Furthermore, we revealed that the M-protein Fab glycoprofiles are mainly complex type glycans that structurally differ significantly from Fc-glycans. Together, glycoproteogenomics opens new opportunities to study *de novo* M-protein Fab N-glycosylation and will contribute to elucidate the mechanisms of M-protein pathogenicity.

## Materials and Methods

### Sequence analysis from public databased to determine M-protein Fab N-glycosylation site

The construction of M-protein Fab region sequences has been described extensively elsewhere^3,6^. In short, RNA-seq datasets from myeloma plasma cells were processed by MiXCR to construct the CDR3 sequences^22^. In case of incomplete sequences, Targeted Assembly of Short Sequence Reads was used^23^. Nucleotide sequences were then translated into (poly)peptide sequences and germline amino acid alterations were determined by HIGH-VQUEST on the IMGT platform.

The Multiple Myeloma Research Foundation (https://themmrf.org/finding-a-cure) provided access to the data from the CoMMpass study on the NIH dbGAP platform (Accession: phs000748.v6.p4). At the time of analysis, 107 full length Fab region sequences were available for analysis.

The cAb-Rep is a database of curated immunoglobulin repertoires from both healthy and diseased donors^24^. The cAb-Rep database was accessed through the web browser interface, where FASTA files containing the nucleotide sequences of the Fab region of immune repertoires of all healthy donors were retrieved. Sequences from either IgG heavy chain (HC) or Kappa or Lambda LCs were used for sequence analysis.

The Fab region sequences were analyzed by an in-house made R script (Version 3.5.3) that determines whether an N-glycosylation motif is present in the sequence. An N-glycosylation motif was defined as a asparagine residue followed by any amino acid besides proline followed by a serine, threonine or cysteine (non-canonical) residue (Asn-X-Ser/Thr/Cys)^25^.

### Samples from patients with Multiple Myeloma

Genomic data and serum samples were collected from the IFM 2009 clinical study (ClinicalTrials.gov identifier NCT01191060 trial) after written informed consent, and clinical and genomic data were de-identified in accordance with the Declaration of Helsinki and approval for this study was provided by our Institutional Review Board (2018-4140)^26^. The Fab M-protein sequences were determined by analysis of a 100 000 read sample extracted from RNA sequencing data. All serum samples were stored at −80 °C upon arrival.

### M-protein purification and digestion

The purification of the M-proteins from serum was performed based on the LC type of the M-proteins. For M-proteins containing a Kappa LC, Capture Select Kappa XP affinity matrix (Thermo Fisher Scientific, Waltham, Massachusetts, USA) were used, and for M-proteins containing a Lambda LC, Capture Select Lambda Hu affinity matrix (Thermo Fisher Scientific) was used. 50 μl of the affinity matrix was transferred to a 0.45 μm 96-wells filter plate (AcroPrep Advance 1 mL, PALL, New York, New York, USA). The affinity matrix was washed three times with 400 μl PBS (Gibco, Waltham, Massachusetts, USA). 200 μg M-protein was used for the purification for most patients, while for patient 7 100 μg was used due to the low M-protein concentration, The M-protein concentrations were determined by serum protein electrophoresis^6^. The corresponding amount of serum was diluted 6-fold in PBS. The diluted serum was incubated with the affinity matrix for 1 hour at room temperature with agitating on a thermomixer at 1000 rpm. The depleted serum was removed by spinning the plate at 1000 g for 1 min at room temperature. The affinity matrix was washed twice with 400 μl PBS supplemented with 0.5 M NaCl (Sigma Aldrich, Saint Louis, Missouri, USA). To elute the M-proteins, the affinity matrix was incubated with 50 μl Glycine (pH=3, Sigma Aldrich) for 5 minutes at room temperature. The eluted M-protein fractions were collected in Eppendorf tubes prefilled with 5 μl 2 M TRIS.HCl (pH=7, Sigma Aldrich), to neutralize the solution. The samples were stored at −80 °C until further processing.

For digestion, 25 μg purified M-protein was used. The M-protein solutions were diluted 1:1 with 8 M Urea (Sigma Aldrich) in Tris.HCl (pH=7) to unfold the proteins. Next, the samples were reduced by addition of 1 μl 10 mM dithiothreitol (Sigma Aldrich) and incubation for 30 minutes at room temperature. Subsequently, the samples were alkylated by addition of 1 μl 50 mM 2-chloroacetamide (Sigma Aldrich) and incubation for 20 minutes at room temperature in the dark. The alkylated samples were diluted 1:4 in 50 mM ammonium bicarbonate (Sigma Aldrich) before addition of a mixture of Trypsin (Promega, Madison, Wisconsin, USA) and LysC (FUJiFILM Wako, Akasaka, Minato, Tokyo, Japan). Both proteases were added in a 1:50 ratio. The samples were incubated overnight at 37 °C. The peptide mixtures were stored at −80 °C until further processing.

### LC-MS/MS analysis of the M-protein peptides and glycopeptides

1 μl of each sample was injected into a nano-HPLC system (nanoElute, Bruker Daltonics, Billerica, Massachusetts, USA) configured in trap set-up. The liquid phase consisted of water (Buffer A, Hipersolv, VWR, Radnor, Pennsylvania, USA) and acetonitrile (Buffer B, ULC/MS grader, Biosolve, Valkenswaard, NL) supplemented with 0.1% formic acid (Biosolve) and 0.01% trifluoroacetic acid (Sigma Aldrich). Samples were loaded onto a trap column (Thermo Fisher Scientific Acclaim PepMap RSLC, 20 mm length, 75 um I.D, 3 um particle size) with 4 times the pick-up volume + 2 μl buffer A at 800.0 bar. The peptides were subsequently separated on the analytical column (Bruker Daltonics, nanoElute FIFTEEN, 150 mm length, 75 μm I.D, 1.9 μm particle size) with a linear gradient from 3 to 42% buffer B in 60.0 min at 500 nL/min. The column was operated at 45 °C. To increase ionization efficiency of hydrophilic compounds such as glycopeptides, a nano-electrospray source (CaptiveSpray, Bruker Daltonics) was used equipped with Nanobooster (Bruker Daltonics) filled with acetonitrile (Biosolve, operated at 0.2 bar). (Glyco)peptides were analyzed on the timsTOF Pro 2 (Bruker Daltonics) operated in PASEF-DDA mode. MS and MS/MS spectra were acquired over 100 to 4000 m/z. The mobility range for the TIMS was set from 0.70 – 2.00 Vs/cm2 with a 100 ms accumulation and ramp time for a 100% duty cycle. Data-dependent acquisition was performed using 8 PASEF MS/MS frames per cycle, with an active exclusion time of 0.4 min or 4-fold increase in intensity. The target intensity was set to 100 000 counts and intensity threshold to 2 500 counts. For the acquisition of MS/MS spectra Collision Induced Dissociation and TIMS stepping was used with a high and low collision energy spectrum. A high collision energy spectrum was acquired at a collision energy ranging from 35.00 eV at 0.60 Vs/cm2 to 131.25 eV at 2.00 Vs/cm2, the Collision RF was set to 1 600 Vpp, transfer time to 64 μs and Pre Pulse Storage time to 10 μs. A low collision energy spectrum was acquired at a collision energy ranging from 20.00 eV at 0.60 Vs/cm2 to 75.00 eV at 2.00 Vs/cm2, the Collision RF was set to 2 000 Vpp, the transfer time to 100 μs and the Pre Pulse Storage time to 13 μs.

### Data Analysis

The Data Analysis of glycoproteomic LC-MS/MS datasets and their integration with genomic data is extensively described in the supplementary information (Supplementary Materials & Methods). In short, LC-MS/MS datasets were analyzed to identify glycopeptide fragmentation spectra, of which the glycan moiety and peptide were identified based on database searches. For the peptide database search, a custom database was created containing the M-protein Fab region sequences. The Mass spectrometry proteomics data have been deposited to the ProteomeXchange Consortium via the PRIDE ^27^ partner repository with the dataset identifier PXD040718.

### 3D modeling of the Fab regions and their N-glycosylation sites

The 3D structures of the M-protein Fab regions were modelled by close homology crystal structures in AbyMod^28^. To acquire the model both a HC and LC Fab sequence has to be known. Since the M-protein of patient 7 consists of a LC only (Table 1) this patient was excluded from analysis. Sequences were entered and the known crystal structures with the highest sequences homology and resolution were selected. 3D structures of the Fab regions were exported and analyzed in pymol (version 2.4.0). The solvent-accessible surface area (SASA) was calculated with the function “get.area”.

**Table 1.**
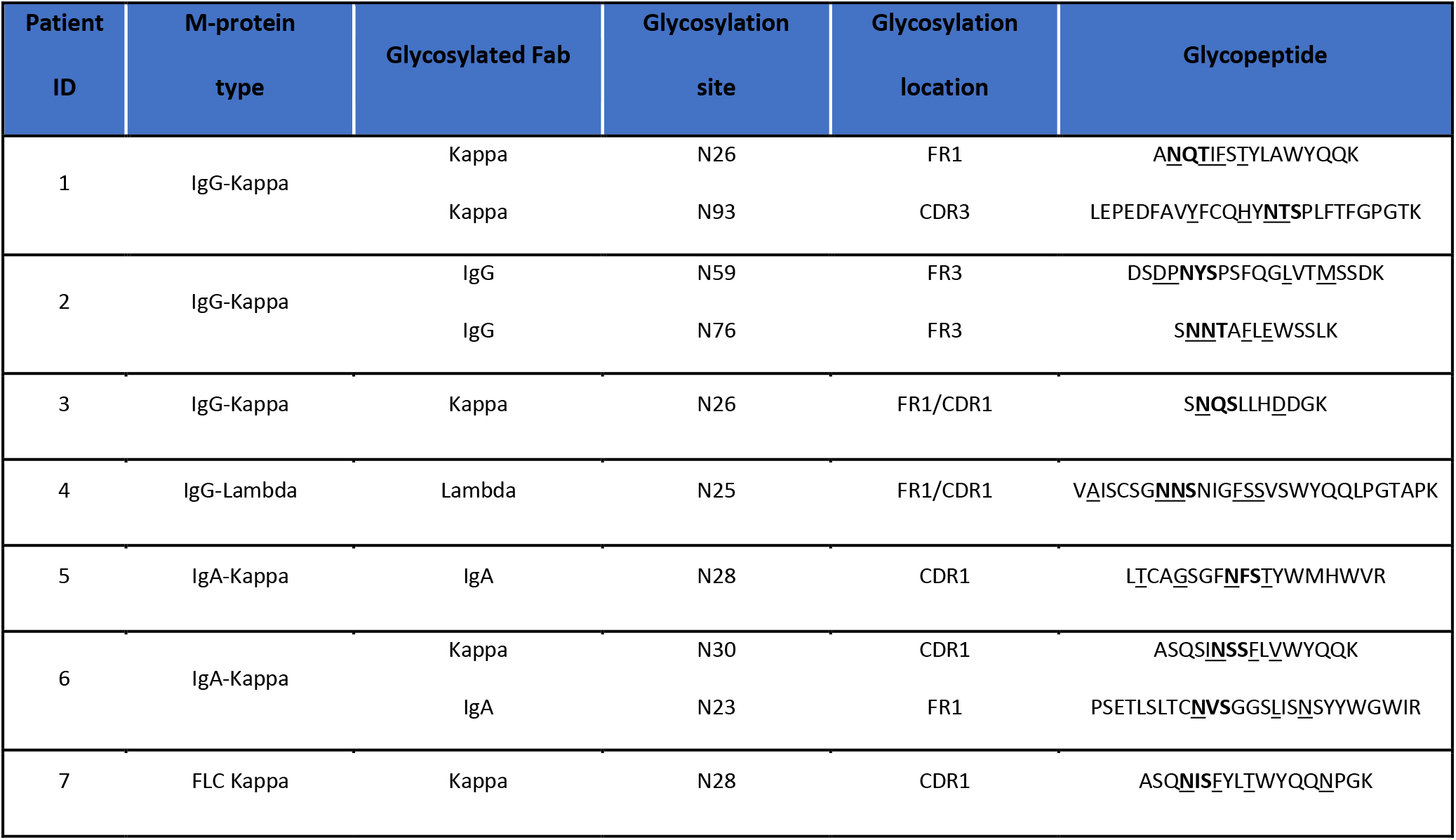
M-protein characteristics and the *de novo* Fab N-glycosylation sites of the patients with MM. Amino acid alterations compared to germline are underlined and N-glycosylation motifs are indicated in bold. FR, Framework; CDR, complementary determining region.

## Results

### Increased *de novo* M-protein N-glycosylation sites in patients with Multiple Myeloma

The presence of Fab N-glycosylation on the M-protein LC has proven prognostic value for plasma cell dyscrasia, mainly in AL amyloidosis^16,17^. Here, we investigated whether Fab N-glycosylation sites are more commonly observed in the M-proteins of patients with Multiple Myeloma compared to immunoglobulins in healthy individuals. To determine the occurrence of an N-glycosylation sites in the M-proteins of patients with MM, we reconstructed the M-protein Fab sequences based on RNA-seq data present in the CoMMpass database. We could reconstruct the full-length sequence of 104 HCs and 93 LCs. An N-glycosylation site was observed in 11.5% of the HCs and in 11.8% of the LCs (Figure 1A). This implies that Fab-glycosylation sites were observed in 22% of the M-proteins. The NxS motif was most dominantly observed in HCs (9 out of 11) and the NxT motif (7 out of 11) was most often seen in LCs. The non-canonical N-glycosylation motif NxC was observed in one LC sequence (Figure 1B). The location of the glycosylation sites was predominantly confined to the region between Framework 1 (FR1) and FR2 in the LC sequences (Figure 1C, red). N-glycosylation sites of the HC were observed over the entire Fab region, with a hot spot at complementary determining region (CDR) 2 and FR3 (Figure 1C, blue).

**Figure 1.**
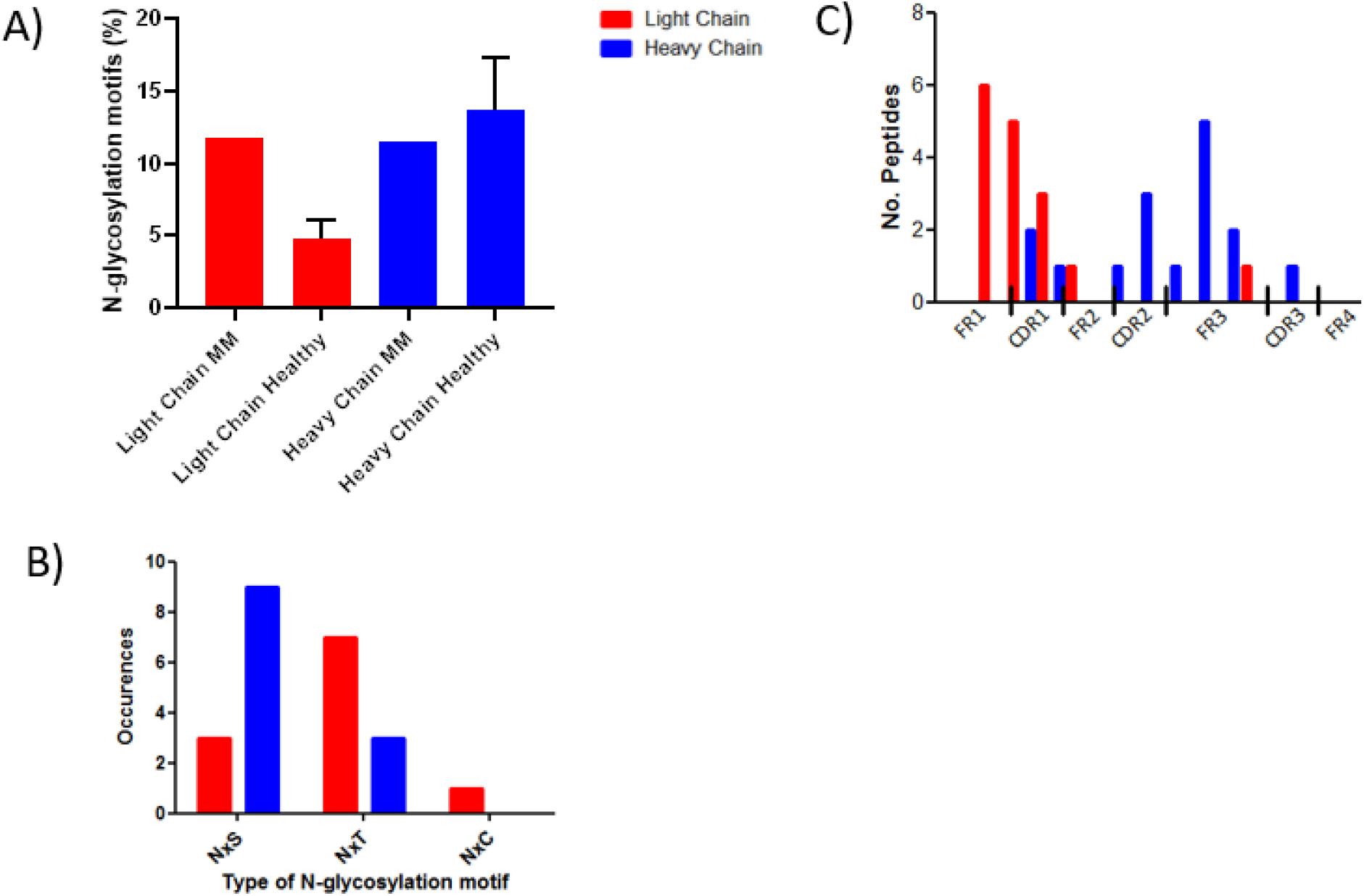
An increase in the occurrence of light chain N-glycosylation sites of the M-protein Fab region is associated with Multiple Myeloma. A) A comparison of the occurrence of Fab N-glycosylation sites (%) on the heavy and light chain of M-protein in patients with Multiple Myeloma and healthy individuals (n=19 for the heavy chains and n=31 for the light chains), reveals that light chain N-glycosylation is more than 2 fold more common in M-proteins than expected from the polyclonal background. NxS N-glycosylation motifs are more common in light chains (red bars) and NxT motifs are more common in heavy chains (blue bars) of M-proteins in patients with MM. C) Glycosylation sites in light chains (red bars) are confined to the Fr1/CDR1 in the V(D)J gene, while in heavy chains (blue bars) the glycosylation sites can be identified across the entire gene. CDR, complementary determining region; FR, Framework; MM, Multiple Myeloma.

Next, we determined the occurrence of Fab regions containing a *de novo* N-glycosylation site in the polyclonal immunoglobulin population of healthy donors available in the cAb-Rep database^24^. This database provided access to sequences of polyclonal HC of 19 healthy individuals (average sequences per donor = 4645) and sequences of polyclonal kappa + lambda LCs of 31 healthy individuals (average sequences per donor = 4809). On average, 13.7% of the HC Fab regions harbored a Fab N-glycosylation site (sd = 3.6 %, range = 4.9 – 21.10 %). On average, 4.8% (sd = 1.3 %, range = 2.2 – 7.4%) of the LCs harbored a Fab N-glycosylation site (Figure 1a), which is significantly lower compared to the HC (p<0.0001). In conclusion, the occurrence of Fab N-glycosylation sites in the HC s of M-proteins in patients with MM was similar in healthy donors (11.5% versus 13.7%, Figure 1A). For the LC a more than 2-fold increase in the occurrence of Fab glycosylation sites in M-proteins compared to healthy donors was observed (11.8% versus 4.8%).

### Studying M-protein *de novo* glycosylation using glycoproteogenomics

Protein N-glycosylation is a non-template driven process capable of generating thousands of unique N-glycan structures for several hundred unique sugar compositions. As such, the collection of N-glycan structures and their stoichiometry that decorate a single N-glycosylation site cannot be derived *in silico* from sequence information. Moreover, the mere presence of an N-glycosylation sequon does not guarantee any glycan occupancy. Therefore, we established a glycoproteogenomics workflow to characterize the glycosylation status of *de novo* N-glycosylations sites of patient-specific genomic alterations.

Firstly, based on RNA-seq datasets from malignant plasma cells in the bone marrow of patients with MM, the sequence of the Fab region of the M-proteins was predicted (Figure 2 Genomics). These Fab protein sequences were used to identify the presence of N-glycosylation sites and *in silico* digestions were performed to predict the theoretical glycopeptides. Secondly, glycoproteomic analysis of the M-proteins was performed (Figure 2, Glycoproteomics). Thirdly, glycoproteomics data were mapped on the RNA-seq derived Fab sequences of the M-proteins to annotate the clonotypic glycopeptides and determine their glycoprofiles (Figure 2, Integration of genomics and glycoproteomics). Furthermore, the Fc region N-glycosylation was also monitored. An example of the generated data is shown for patient 1 (Figure 2b), in whom we identified two N-glycosylation sites in the Fab sequence (N15 and N128). We were able to identify site specific N-glycan profiles and differentiate them from Fc region N-glycosylation. Next, we applied this workflow to a cohort of 41 patients with MM.

**Figure 2.**
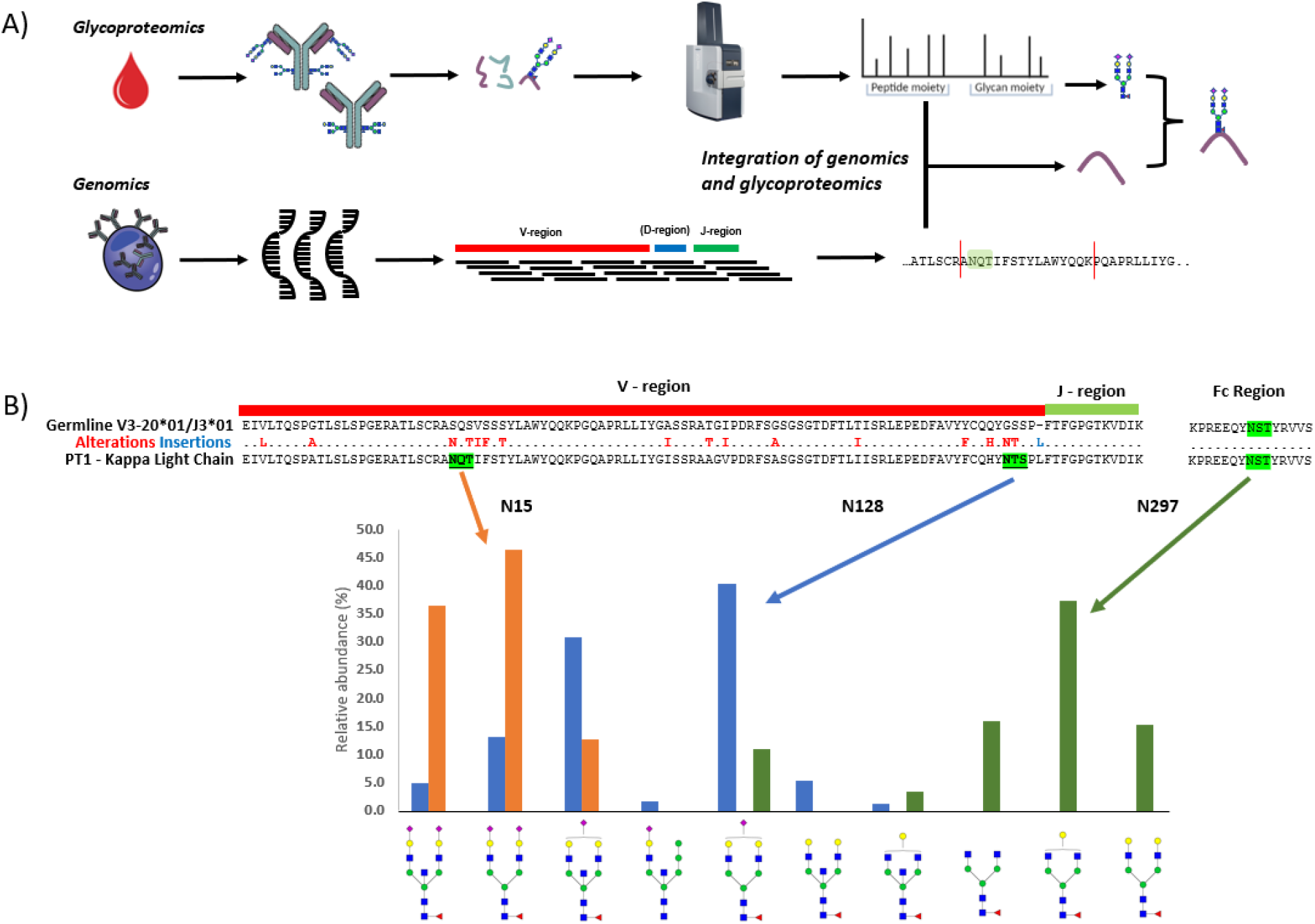
Glycoproteogenomics, the integration of genomics and glycoproteomics to study *de novo* Fab N-glycosylation of M-protein. A) Schematic overview of the glycoproteogenomics workflow where genomics is integrated with LC-MS/MS based glycoproteomics to study *de novo* Fab N-glycosylation. B) An example of generated data for a patient, whereby, based on genomic analysis, two *de novo* N-glycosylation sites were predicted (highlighted in green). Subsequent glycoproteomics analysis revealed two distinct glycoprofiles (orange and blue) on the two glycosylation sites. In parallel, the glycoprofile of the Fc was also detected (darkgreen), showing a different glycoprofile compared to the Fab region.

### Profiling *de novo* N-glycosylation sites in the Fab regions of M-proteins

Based on the RNA-seq derived Fab region sequences, we identified 10 N-glycosylation sites in 7 out of 41 patients with MM (Table 1). 9 out of 10 sites were the result of amino acid alterations and were thus *de novo* introduced into the Fab region. Only the IgG N59 site in patient 2 was already present in the germline sequence (IGHV5-10-1*04). Fab glycosylation sites were present on both the LC (n=6) and HC (n=4) of the M-protein. For patient 1 and 2, two Fab glycosylation sites were identified in a single chain, while for patient 6 a glycosylation site in both the LC and HC was present. In Table 1 each patient- and myeloma specific glycopeptide (clonotypic glycopeptide) is shown.

For the 7 out of 10 of the Fab glycosylation sites we obtained glycan signals for all matching clonotypic glycopeptides, indicating that these sites were 100% occupied by glycans or near 100% due to peptide specific sensitivities (Figure 3A). For two sites, we observed partial glycan occupancy (51% for the Fab site Kappa N128 from patient 1 and 94% for Kappa N30 from patient 6), while the IgG N59 Fab site from patient 2, present in the germline sequence, was completely unoccupied.

**Figure 3.**
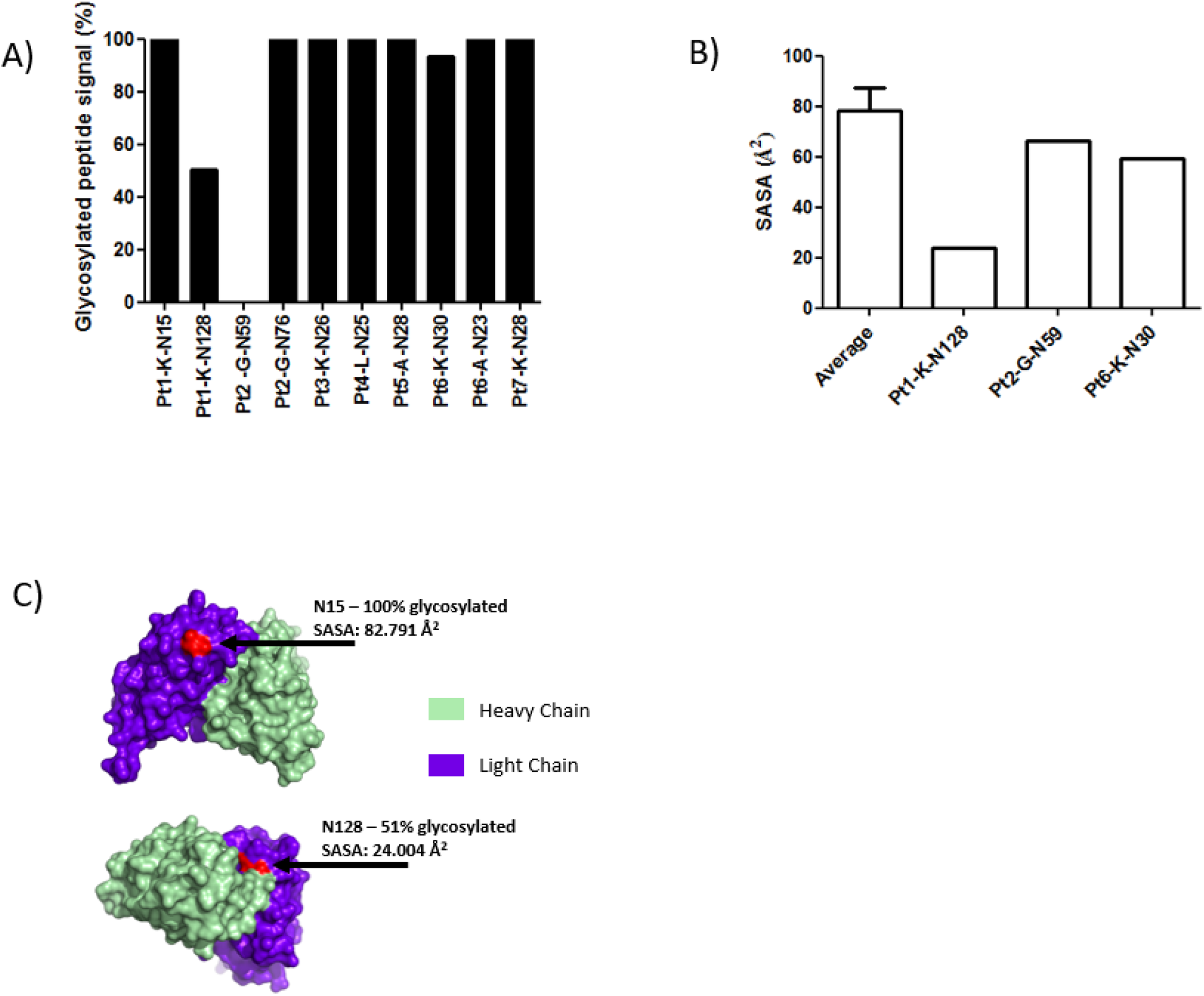
Occupancy of *de novo* Fab N-glycosylation sites on the Fab region of M-proteins. A) The relative signal of the measured glycosylated peptide signal per Fab N-glycosylation reveals three sites with a less than 100% glycan occupancy. Note that Pt2_G-N59 is an N-glycosylation site encoded in the germline. B) The surface accessibility (SASA) was calculated based on 3D modeling of the Fab regions. For the three sites with a less than 100% occupancy the SASA is depicted, as well as the average of the SASA of the fully occupied sites (n=7). Only for site N128 from patient 1 a decrease in SASA was observed. C) 3D structures of the Glycosylation sites on the M-protein of patient 1 unravels the buried position of N128. In green the heavy chain, and in purple the light chain is depicted.

The amino acid sequence context of the N-glycosylation site can influence the glycan occupancy^25^. Classically, a proline between the asparagine and serine/threonine residue prevents N-glycosylation. Similarly, a proline residue succeeding the serine or threonine residue (NxS/TP) can also hamper N-glycosylation^25^. The succeeding proline in the glycosites of patient 1 (NTSP) and the pre- and succeeding proline in patient 2 (PNYSP) may thus explain the (partial) N-glycosylation observed in these M-proteins (Table 1). The loss of glycosylation observed at the Fab glycosylation site in patient 6 cannot be explained by the presence of a proline in the sequence. Surface accessibility of the asparagine residue for N-glycosylation enzymes might be of importance. To study this, we determined the surface accessibility (SASA) for the corresponding asparagine residues within the M-proteins. 3D modeling of the Fab region of each M-protein showed that the SASA of 24.004 Å2 in Pt1-K-N128 (51% glycosylated clonotypic glycopeptide signal) was significantly lower compared to the average SASAs present on the other N-glycosylation sites (Figure 3B). In the 3D model it is apparent that the asparagine is hidden in between the LC and HC (Figure 3C). The low SASA indicates a decreased accessibility for protein glycosylation enzymes and could explain the low site occupancy observed on Pt1-K-N128.

Together, these data highlight that the genomics and glycoproteomics ought to be combined in order to decipher *de novo* N-glycosylation status on M-proteins.

### Comparison of M-protein Fab and Fc region N-glycosylation

We further characterized the glycoprofiles of M-proteins from patients with MM. The Heavy Chain Fab region and Light Chain Fab region are decorated with fairly similar glycans (Figure 4A yellow and pink shade). These are mainly complex type, highly sialylated and fully fucosylated glycans with various degrees of bisecting structures, such as H5N5S1F1 and H5N4S2F1. Next, we compared the glycans observed on the Fab region with the ones on the Fc regions. We confirmed that N-glycosylation occurred on the Fc region of both IgA (N459 and N263, Figure 4A green shades) and IgG (N297, Figure 4A blue shades) M-proteins. Similar to the Fab region, the Fc regions are also mainly decorated with complex type glycans. High mannose glycans were exclusively observed on the Fc region of IgA. Even though the major structural class is similar between Fab and Fc region glycans, within this class a clear distinction between the glycans observed on the two regions can be made. The Fc region glycans are for the majority simpler glycans containing either only fucose (H3N4F1 IgG Fc region) or sialic acid (H5N4S1 on IgA) compared to the fully decorated glycans observed on the Fab of M-proteins.

**Figure 4.**
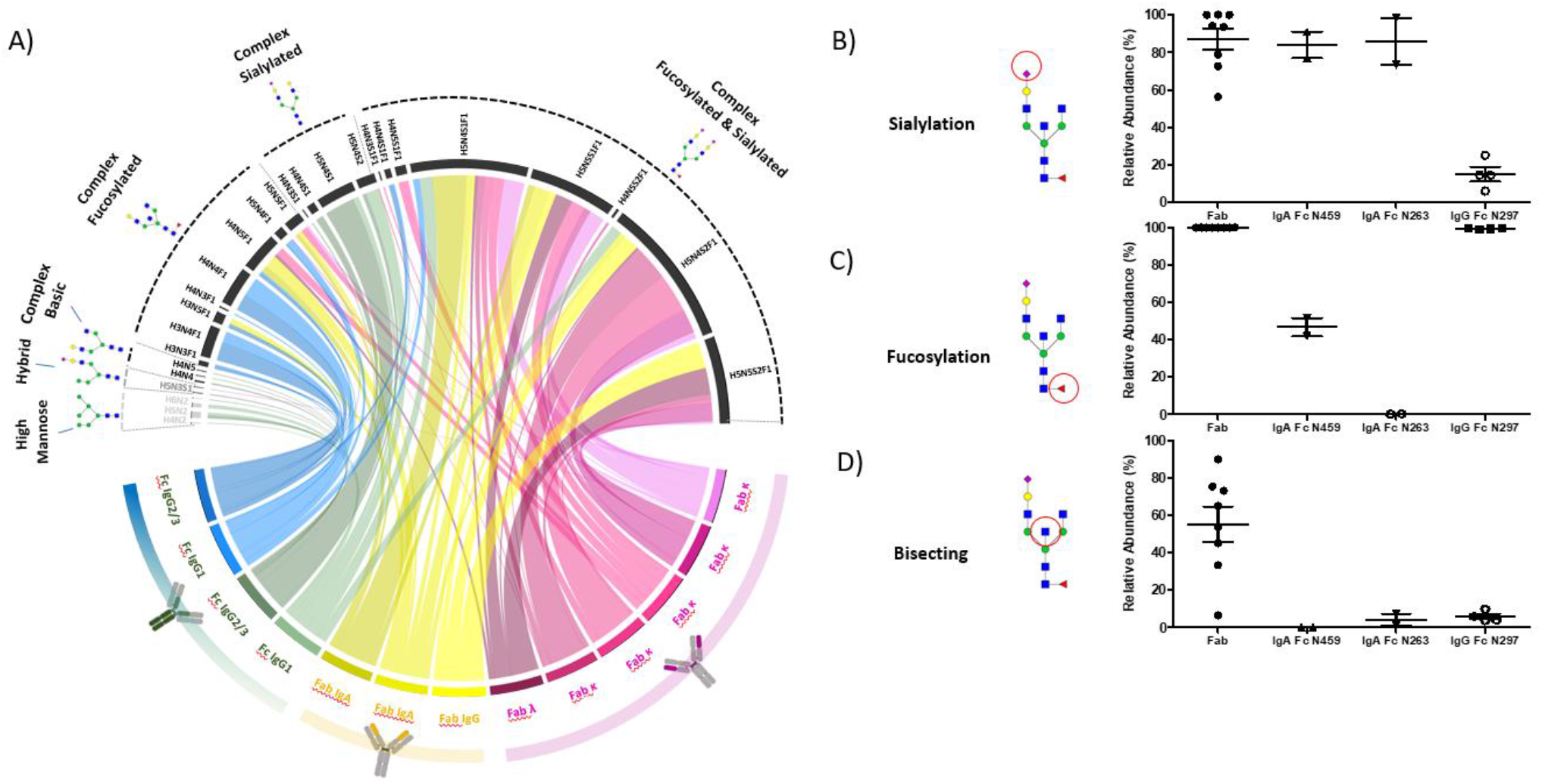
Glycan types and structures differ between Fab and Fc N-glycosylation. A) cord plot connecting the N-glycosylation sites on the bottom with observed N-glycans on the top (>1% abundance) reveals unique profiles that distinguish Fc glycans and Fab glycans. The cord size represents relative abundances and the colors are grouped based on the chain of the N-glycosylation site. B) the sialylation levels of Fab glycans were higher compared to IgG Fc glycans, while C) fucosylation levels were lower in IgA Fc region glycans. D) bisecting structures were more abundant in Fab region glycans compared to Fc regions.

Next, we grouped and quantified the observed glycans based on structural elements; sialylation, fucosylation and bisecting GlcNaC containing structures. The level of sialylated glycans was comparable between the Fab and IgA Fc region (Figure 4B). The sialylation of glycans in the IgG constant region was significantly lower (p <0.0001). The fucosylation levels were comparable between Fab and IgG Fc regions, but were significantly lower for Fc IgA, with an almost complete absence of fucosylation on site N263 (Figure 4C). Finally, a striking difference was observed regarding the abundancy of bisecting structures in the Fab glycans compared to an almost complete absence of bisecting Fc glycans (Figure 4D). These data indicate site-specific glycoform structures with significant differences between M-protein Fc and Fab N-glycosylation.

## Discussion

Malignant plasma cells in MM produce and secrete a characteristic monoclonal immunoglobulin (M-protein) that contains a unique Fab region. The uniqueness of the Fab region arises due to V(D)J rearrangements and somatic hypermutations. This can lead to the introduction of myeloma- and patient-specific *de novo* N-glycosylation sites. Recently, evidence for the prognostic value of Fab N-glycosylation of the M-protein in certain monoclonal gammopathies was provided ^16,17^. However, methods to characterize these N-glycosylation sites thus far lacked specificity to determine the occupancy and glycoforms of both the HC and LC in a site-specific manner^13,14,16^. Here, we present a glycoproteogenomics approach that integrates genomics with glycoproteomics, which we applied to characterize *de novo* Fab N-glycosylation on the M-proteins of patients with MM in unprecedented detail. Acquiring in-depth information on the exact M-protein glycoprofiles might provide a biomarker to predict disease progression or contribute to our understanding of M-protein pathogenicity.

The strength of the combination of -omics technologies became apparent when we applied the glycoproteogenomics approach to a cohort of 41 patients with MM. Ten Fab N-glycosylation sites on the M-protein of seven patients were predicted based on genomic data. Subsequent glycoproteomic analysis confirmed glycan occupancy in nine of these sites, for the first time we were able to characterize multiple N-glycosylation sites on the same Fab and study their glycoform profiles separately. Two Fab glycosylation sites were partially occupied (51% and 94% for Patient 1 N128 and patient 6 N30, respectively). One site, already present in germline, appeared to be fully unoccupied. Based on the amino acid sequence information of the Fab regions, we could not predict the degree of glycan-occupancy of a specific site. The fully unoccupied site was the only Fab N-glycosylation site present in the germline sequence (IGHV5-10-1). Relatively few immunoglobulin Fab alleles contain such a conserved N-glycosylation site (IGHV1-8, IGHV4-34, IGHV5-10-1, IGLV3-12, and IGLV5-37)^29^; due to somatic hypermutations a proline residue is present in the N-glycosylation sequon preceding the asparagine residue. We hypothesize that during antigen binding affinity selection there was an advantage for this non-glycosylated version of the B cell receptor.

The glycans observed on the Fabs of the M-proteins of patients with MM were almost all of the complex type with a high degree of sialylation, fucosylation and bisecting GlcNAc containing structures. The M-protein Fab glycan structures were distinguishable from the glycans on the Fc region of the M-protein in terms of sialylation (low degree of sialylation in Fc IgG), fucosylation (low degree of fucosylation in Fc IgA), and both the Fc IgG and Fc IgA had a lower degree of bisecting glycans. It is hypothesized that the limited Fc sialylation in IgG antibodies is due to the partially shielded inner facing position of the glycans^12^.

Even though the N-glycosylation sites on the Fab regions were for the majority occupied by complex type glycan structures, we did observe glycan heterogeneity between patients, between chains and even within the same Fab chain of one M-protein. The glycan heterogeneity was caused by the wide range of sialic acid content (56% - 100%) and bisecting GlcNac containing species (6%-90%). The glycan variation was much less pronounced on the conserved Fc region glycans. This could suggest that the glycoforms on the Fab region are less regulated. Van de Bovenkamp et al. hypothesized that Fab glycans may regulate antigen binding^13^. The variation in Fab glycoforms could thus indicate selection based on the highest affinity for the antigen. In contrast to our findings, complex and high mannose Fab glycan structures have also been described on both Follicular Lymphoma M-proteins^30^. These structures seem unlikely due to the accessibility of the N-glycosylation sites, which allows for full maturation of these sites.

There was a more than two-fold increase in the occurrence of Fab N-glycosylation sites in the M-protein LCs of patients with MM (11.8%) compared to polyclonal antibodies in healthy individuals (4.8%), indicating an association between MM and an increase in M-protein LC Fab N-glycosylation. This phenomenon was not seen on the M-protein HC, as in agreement with earlier studies^31^. For M-proteins from patients with monoclonal gammopathy of undetermined significance, a benign monoclonal gammopathy, it was shown that the LC Fab N-glycosylation occupancy is 6%^17^. This is similar to the occupancy we observed in healthy controls. In AL-amyloidosis, a LC plasma cell malignancy, the occurrence of monoclonal LC N-glycosylation was also increased (~16%)^16^. In AL amyloidosis the Fab LC glycans contribute to the pathogenicity of the M-proteins^32^. M-protein pathogenicity in MM is poorly understood^33,34^. Based on both mice-models and human data, Westhrin et al. showed that abnormal M-protein Fc glycosylation may promote bone loss in MM and altering IgG glycosylation may be a therapeutic strategy to reduce bone loss^35^. Thus far, no causal relationship between Fab N-glycosylation and M-protein pathogenicity has been described in MM, however it is speculated that Fab N-glycosylation might contribute to M-protein pathogenicity^34^. Recent data strongly suggests that the presence of M-protein Fab N-glycosylation is a risk factor for progression from MGUS to AL-amyloidosis and other plasma cell disorders^17^. Methods to study this have relied on the discrimination between glycosylated and non-glycosylated M-protein LCs. Even though there are ongoing efforts to optimize these techniques to also elucidate glycan classes and profiles, the level of insight glycoproteogenomics offers is unprecedented^36^. We hypothesize that the additional information about glycosylation of the Fab region of the HC, the specific glycan profiles and the exact site on the M-protein Fab, obtained by the unprecedented detail of glycoproteogenomics, will aid in unraveling the clinical role of M-protein Fab glycosylation. Further studies in larger, independent cohorts are warranted to study this, not only in MM but in other plasma cell dyscrasias as well.

Proteogenomics, the integration of genomics and proteomics, is an emerging technological development in the multi-omics field and has the potential to improve cancer characterization, diagnosis, and even advance cancer therapy(^37–39^). Furthermore, such integration efforts are also being used to sequence antibody repertoires by MS^40^. With glycoproteogenomics, we can now relate genomic alterations to detailed glycosylation phenotypes in a clinical setting. In the current study, glycoproteogenomics is applied to study *de novo* Fab N-glycosylation, but we believe that this can also contribute to other fields. Acquisition of holistic genomic data of patients is becoming more custom in clinical practice. In such genomic data the deletion or acquisition of glycosylation sites through mutations/truncations can be distilled, followed by subsequent glycoproteomics analysis to dissect the glycosylation phenotype and whether this has clinical consequences for the patients. The glycoproteogenomics workflow introduces a novel level of personalized diagnostics and can aid in improving patient care.

## Acknowledgements

This work was financially supported by two grants from the Dutch Cancer Society for JFMJ (#10817 and #14465) and by Sebia (Lisses, France). This research was supported by ZonMw Medium Investment Grant (40-00506-98-9001) and was part of the Netherlands X-omics Initiative supported by collaboration project “EnFORCE” (LSHM21032) which is co-funded by the PPP Allowance made available by Health~Holland, Top Sector Life Sciences & Health, to stimulate public-private partnerships. The authors further acknowledge the Multiple Myeloma Research Foundation for providing access to the CoMMpass study data through NIHdbGap.

## Author contributions

PL performed experiments, data analysis and writing of the manuscript. MB/PK performed data analysis and contributed to writing of the manuscript. SNa, CW, MP, SNo, MD, IJ, AG, JG contributed to writing the manuscript. TD provided samples from the global biobank and contributed to writing. DL helped conceptualize the study and contributed to writing. HW/JJ conceptualized the study, contributed to data analysis and writing of the manuscript.

