## Supplementary Materials&Methods for "*N*-linked glycosylation of the M-protein variable region: Glycoproteogenomics reveals a new layer of personalized complexity in multiple myeloma"

**Supplementary Information**

This file contains supplementary Materials & Methods

**Supplementary Materials & Methods.**

**Data Analysis of glycoproteogenomics data set**

LC-MS/MS datasets were converted to mascot generic files (MGF) with custom made scripts in DataAnalysis (Bruker Daltonics, v 5.0). The processing of MGF’s was performed in BioPharma Commpass (BPC, Bruker Daltonics, version 3.0.0.19). Datafiles were uploaded and classification of glycopeptides fragmentation spectra was performed. The spectra were classified as glycopeptide spectra if at least 2 consecutive m/z distances (tolerance of 0.05 Da) consistent with the tetrasaccharide core of an N-glycan were observed (Peptide-HexNAc-HexNAc-Hex-Hex). For each glycopeptide spectrum the peptide and glycan mass were calculated using N-glycan core fragmentation pattern matching, for subsequent identification of each respective moiety by protein- and glycan-database search engines. To identify the peptide sequences, a database search of the peptide and glycopeptide classified spectra was performed. Spectra were submitted from BPC3 to an in-house mascot server (v2.5, Matrix Science). Here, the spectra were searched against a custom made database consisting of the uniprot human proteome database supplemented with the RNA-seq derived M-protein Fab region sequences to allow the identification of the (glyco)peptides from this region. Percolator was used to achieve 1% FDR and peptide spectrum matches were accepted if the mascot score was above the percolator adjusted FDR score threshold (>10). To identify the N-glycans attached to the glycopeptides, the glycan mass and corresponding fragments in the glycopeptide classified spectra were analyzed by GlycoQuest (BioPharma compass version 3.0.0.19). Here, the database from the Consortium for Functional Glycomics was used. The MS tolerance was set to 10 ppm and MS/MS tolerance to 0.05 Da. The score, fragmentation coverage (%) and intensity coverage (%) had to be higher than 10 for a glycan assignment. Subsequently the peptide and N-glycan search results were merged to obtain glycopeptides identifications. In case of partially elucidated glycopeptides the respective peptide- or glycan-moiety were annotated using co-eluting fully elucidated glycoform identifications if the calculated respective moiety mass was within +/- 20ppm mass tolerance as described in Wessels et al^1^. Based on the (glyco)peptides identifications a target list was constructed of the glycopeptides derived from the M-protein Fab region and immunoglobulin constant region, containing monoisotopic *m/z*, *z* and retention time (Supplementary table 1). This list was used to determine the extracted ion chromatogram peak areas of the corresponding targets in the LC-MS/MS dataset. The analysis was performed in Data Analysis (Bruker Daltonics, v5.0) using custom made scripts. Here, the targets were extracted within a 1.5 min retention time window and 0.1 m/z tolerance. The mass differences in the isotope pattern were checked to belong to the corresponding charge within +/- 20 ppm tolerance. Finally, the chromatographic signal of the mono- up to the 4th isotopic signal was summed and integrated and this area was reported. Occupancy was indicated by comparing the area of non-modified glycopeptides to the sum of the glycan-modified glycopeptides.
